# STOMP-seq: early multiplexing for high-throughput SMART RNA-sequencing

**DOI:** 10.1101/2025.03.28.645277

**Authors:** Matthias Eder, Nicholas Stroustrup

## Abstract

RNA-sequencing provides high-dimensional, quantitative measurements of the states of cells, tissues, organs, and whole organisms. Plate-based RNA-seq protocols allow for a wider range of experimental designs than droplet sequencing methods, but are less scalable due to the practical challenges of plate-based liquid handling. Here, we present STOMP-seq, a method that extends SMART RNA-seq protocols, like Smart-seq2 and Smart-seq3, to include sample-identifying barcodes on the 5’ end of each amplified transcript. These barcodes allow samples to be pooled immediately after reverse transcription, enabling a 12-fold multiplexing strategy that reduces liquid handling complexity and enzyme costs several-fold. Suitable for both manual and robotic library preparation approaches, STOMP-seq reduces protocol execution times four-fold while improving library complexity and coverage. Together, these advantages combine to make possible new large-scale experimental designs, in particular population-scale sequencing projects like the multi-generational study of gene-expression heritability presented here. STOMP-seq offers a “drop-in” replacement for Smart-seq2 and Smart-seq3, removing practical barriers that currently limit the quality and scope of plate-based transcriptomic data.

**Graphical Abstract:** 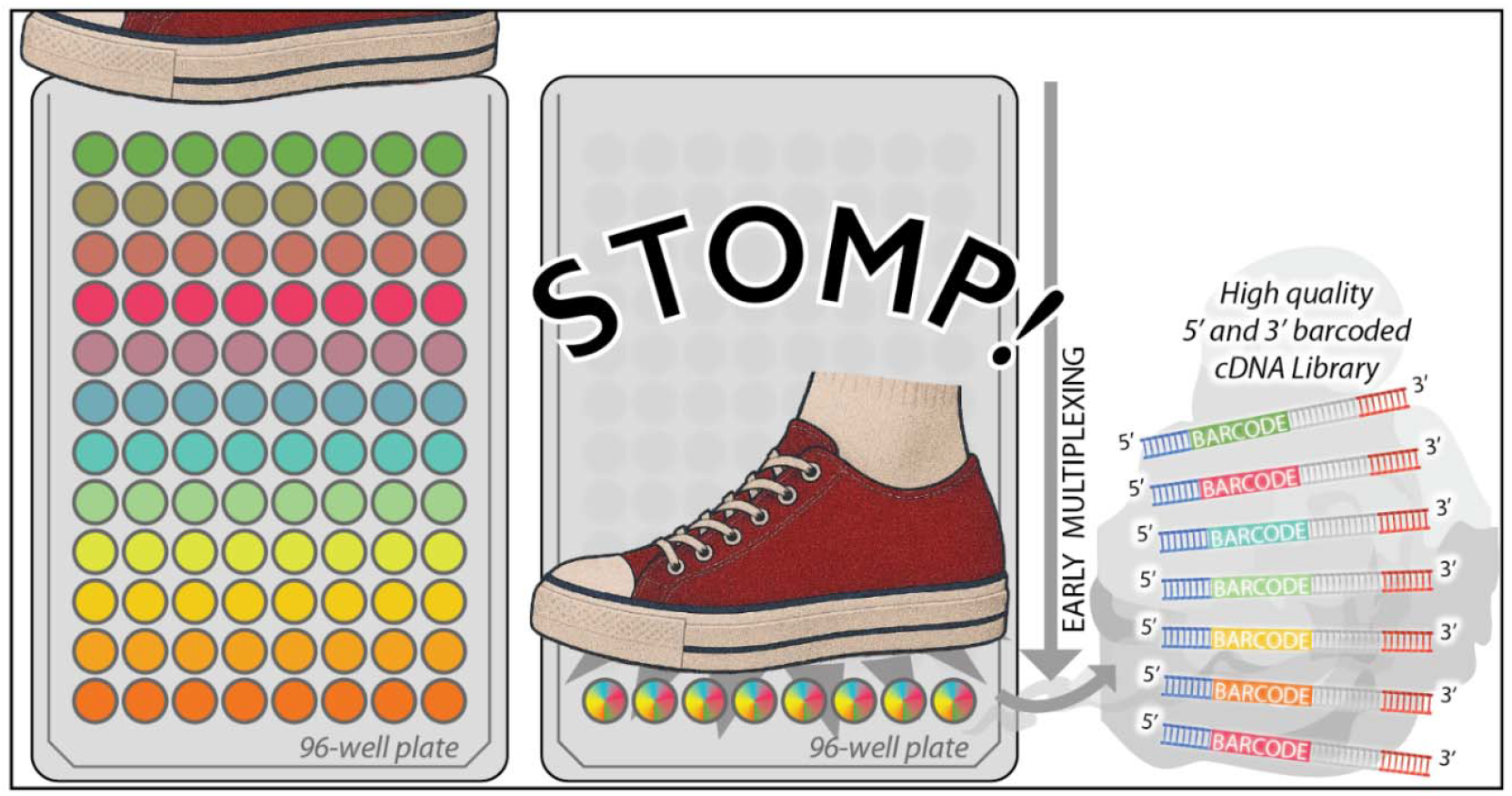

## Introduction

RNA-sequencing provides researchers with rich, multidimensional measurements of the state of cells, tissues, organs, and whole organisms. Compared to droplet or in-situ-based RNA-seq, plate-based methods like Smart-seq2 ^1^ support a wider range of experimental designs, in particular those involving flow cytometry, while producing high-quality results at a price point accessible for single-laboratory, exploratory studies. Recent innovations to SMART (**S**witching-**M**echanism-**A**t-5’-end of **R**NA **T**emplate) family protocols include the addition of UMIs ^2^ and the merging of protocol steps ^3^ to speed up liquid handling in the protocol. We reasoned that an early-pooling, sample multiplexing strategy ^4–6^ could provide substantially greater efficiency gains by reducing several-fold the number of liquid transfers required for traditional SMART protocols.

To enable early pooling, barcodes must be added using initial reverse-transcription (RT) primers, either at the 5’ or 3’ end of cDNA molecules. The 5’-end is a more efficient location for barcodes compared to the 3’ end which necessarily incorporates non-informative poly-dT sequences into the final sequencing libraries ^7^. To enable 5’ barcoding, we developed a new strategy we call STOMP-seq (<u>**S**</u>MART <u>**T**</u>S**O** <u>**M**</u>ultiplexed 5-<u>**P**</u>rime sequencing) that introduces unique, sample-identifying barcodes onto the SMART 5’-end template switching oligo (TSO). During tagmentation, our protocol’s 5’ barcodes additionally reduce the need for large sets of unique double indexing primers, removing a major expense that often blocks utilization of more cost-efficient high-capacity sequencing flow cells.

## Materials and Methods

### Reagents

| REAGENT | SOURCE | IDENTIFIER |
| --- | --- | --- |
| Betaine, 5M | Sigma-Aldrich | Cat# B0300 |
| Dimethyl sulfoxide | Sigma-Aldrich | Cat# D8418 |
| DTT, 0.1M | Invitrogen | Cat# Y00147 |
| ERCC spike-in mix | Invitrogen | Cat# 4456740 |
| GTP | Thermo Fisher Scientific | Cat# R1461 |
| KAPA HiFi Hotstart ReadyMix | Roche | Cat# KK2602 |
| Magnesium Chloride, 1M | Sigma-Aldrich | Cat# M1028 |
| Maxima H-minus reverse transcriptase | Thermo Fisher Scientific | Cat# EP0752 |
| PEG 8000, 50% solution | Sigma-Aldrich | Cat# 81268 |
| Proteinase K | NEB | Cat# P8107S |
| RNasin | Promega | Cat# N2515 |
| Sodium Chloride, 1M | Sigma-Aldrich | Cat# 28-2300 |
| SPRI paramagnetic beads | CRG Genomics Unit | N/A |
| Superscript II reverse transcriptase Kit | Invitrogen | Cat# 18064014 |
| Tn5 transposase | CRG Protein Technologies Unit | N/A |
| Tris-HCL, 1M pH 8.3 | bioWORLD | Cat# 42020414 |
| Triton X-100 | bioWORLD | Cat# 42030000 |
| Tween 20 | bioWORLD | Cat# 42030016 |
| Critical commercial assays |  |  |
| Quant-iT dsDNA Broad-Range (BR) Assay Kit | Invitrogen | Cat# Q33130 |
| Tapestation High Sensitivity D5000 Reagents | Agilent | Cat# 5067-5593 |
| Tapestation High Sensitivity D5000 ScreenTape | Agilent | Cat# 5067-5592 |

#### mRNA Reverse Transcription

To decrease sample preparation time and sequencing cost, we adapted key components of RNA sequencing protocols of the SMART family to allow 5’ sample barcoding and early pooling. For Smart-seq2-based STOMP-seq, cDNA libraries were generated from 4 µL of raw lysate, which was incubated with 4 µL of Smart-seq2 annealing mix ^1^ for 3 min at 72 °C and then immediately put on ice. 12 µL of reverse transcription (RT) mix were then added, containing the same enzyme and buffer composition as the Smart-seq2 RT mix combined with the STOMP-seq barcoded TSOs. Samples were processed using the standard Smart-seq2 RT thermocycler protocol, after which first strand cDNA samples were immediately pooled together. For PCR amplifications, 20 µL of each pool were mixed with 30 µL of IS PCR mix containing the same enzyme and buffer composition as the Smart-seq2 IS PCR amplification mix along with 83.3 nM of both the Smart-seq2 IS primer and the STOMP-seq IS primer.

To additionally adapt the Smart-seq3 protocol for STOMP-seq multiplexing, we generated cDNA libraries from 4 µL of raw lysate, which was incubated with 2 µL of Smart-seq3 annealing mix ^8^ for 3 min at 72 °C and then immediately put on ice. 2 µL of reverse transcription (RT) mix were then added, containing the same enzyme and buffer composition as the Smart-seq3 RT mix combined with the STOMP-seq barcoded TSOs. Samples were processed using the standard Smart-seq3 RT thermocycler protocol, after which first strand cDNA samples were immediately pooled together. For PCR amplifications, 8 µL of each pool were mixed with 12 µL of PCR amplification mix containing the same enzyme and buffer composition as the Smart-seq3 PCR amplification mix along with 83.3 nM of the STOMP-seq IS primer and 16.7 nM of the Smart-seq3 reverse PCR primer.

For both protocols, amplified cDNA was purified using SPRI paramagnetic beads, produced locally to replace AMPure XP beads (Beckman Coulter), using a bead to sample ratio of 0.8. The cDNA library size distributions were measured using a Tapestation 4150 (Agilent) and the overall library concentration was measured using the Quant-it DNA determination kit (Invitrogen) on a plate reader (Tecan).

#### cDNA Tagmentation

To ensure that 5’ barcodes are propagated into final sequencing libraries, we needed to modify the cDNA tagmentation step. By incorporating a TruSeq sequencing initiation sequence - usually ligated during the TruSeq sequencing library preparation - onto the 5’ TSO, we gain the ability to selectively amplify the 5’-end of each full-length cDNA using PCR primers complementary to this TruSeq fragment and the 3’ indexing Nextera primer sequence added during tagmentation— ensuring 5’ barcode propagation. Strategies combining TruSeq and Nextera primers have been proposed previously ^4,6^, but by incorporating the TruSeq “read2” sequence specifically at the barcoded 5’-end, our strategy avoids low sequence diversity (complexity) in “read1” that otherwise decreases sequencing fidelity ^9^. cDNA libraries were tagmented using the Nextera library preparation protocol (Illumina) ^10^. Fragments containing barcoded 5’ ends were selectively amplified using STOMP-seq N7 indexing primers (Tab. S4) complementary to the incomplete TruSeq sequencing initiation sequence added by the STOMP-seq TSO, and Nextera S5 indexing primers. The first gap-filling step of the Nextera library amplification PCR was omitted to suppress generation of S5-S5 fragments (that cannot be sequenced in standard Illumina flow cells) while preserving N7-S5 fragments which contain one complete strand. Amplified sequencing libraries were purified twice using SPRI paramagnetic beads produced locally to replace AMPure XP beads (Beckman Coulter), using a bead to sample ratio of 0.9. Sequencing library size distributions were measured using a Tapestation 4150 (Agilent) and the overall library concentration was measured using the Quant-it DNA determination kit (Invitrogen) on a plate reader (Tecan). Sequencing libraries were pooled in equal masses and library pools were sequenced on either Illumina NextSeq500, NextSeq2000 or NovaSeq6000 machines, with paired end reads of length 38 bp, 52 bp and 52 bp respectively.

#### Gene-expression analysis

FASTQ sequence files from samples processed using STOMP-seq were demultiplexed using fastq-multx version 1.4.2 ^11^, removing both STOMP-seq barcode sequences and GGG overhangs. Demultiplexed FASTQ files were aligned to the reference genome, *C. elegans* Wormbase release WS265, modified to include ERCC spike-ins, using *STAR* version 2.6.0c ^12^. Genome coverage of reads was computed using *BEDTools* version 2.29.1 ^13^. Gene counts were quantified from alignments using *featureCount*s version 2.0.0 ^14^. Transcripts were filtered for lowly expressed genes using *filterByExpr* from the *R/edgeR* package version 4.2.2 ^15^, except for the transgenerational study where a detection threshold of 5 counts in at least 75% of the samples was applied. Sample counts were normalized using normalization factors estimated from the filtered count matrix using *scran* version 1.32.0 ^16^. Differential gene-expression analysis was performed using the DESeq2 package version 1.44.0 ^17^ in R.

### Statistical Analyses

*PCA:* Principal component analysis was performed in R, running *prcomp* on log-transformed counts with 0.5 pseudo counts added. Counts were centered and scaled. Hierarchical clustering of samples was performed using Spearman’s correlation and the Ward algorithm. Benjamini-Hochberg multiple testing correction for p-values was performed.

#### Multigenerational gene-expression comparisons

To identify any parental influence on progeny gene-expression, a regression analysis was performed to identify genes whose expression was more similar between siblings than between cousins. We considered the log-linear regression model 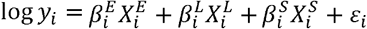 with *y*_*i*_ as the expression of gene *i*, two vectors 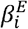 and 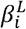 estimating the effect of batch effects of environmental factors and RNA-sequencing library preparation introduced by batches 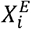 and 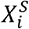 respectively. A final categorical variable 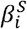 was included which represented the shared influence of a parent across all of its progeny (*i*.*e*., among all members of each sibling group 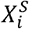). Thus formulated, the regression can identify parental influences common to all siblings through values of 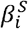 that achieve statistically significant differences between sibling groups 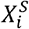. Model parameters were estimated separately for each gene *i* using maximum likelihood estimation using the *glm* package in *R*.

Partial r-squared estimates comparing the influence of batch effects and parental contributions were obtained by comparing the full model to the partial models 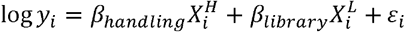 and 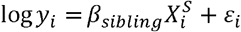 respectively using the *rsq* package in *R*. Sibling group expressions, normalized to exclude batch effects, as plotted in Fig. 5d, were calculated from the full model’s residuals as 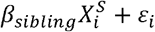.

Parental genes whose expression predict the expression of any gene in their progeny were then identified using partial least squares (PLS) regression. Latent variables were identified by decomposing the matrix of parental gene expression *y* into a set of 5 components, and these variables were then used to predict the expression of a single gene’s expression among progeny. Each gene’s expression in progeny was then analyzed using a separate PLS regression using the *pls* package in *R*. The magnitude of PLS loadings obtained from analyzing shuffled data was estimated by first assigning all individuals in a sibling group to a random parent and then re-running PLS on data permutated to match these new parent-sibling pairings. Hierarchical clustering of parental and progeny gene-expression was performed using *hclust* in *R*.

*R* scripts were run on *R* version 4.1 and RStudio version 2024.09.0.

### Biological Resources

mRNA was obtained from two strains of live *C. elegans* nematodes: QZ0 (Bristol N2), and CB4037 which contains the *glp-1(e2141*) allele responsible for a temperature-sensitive germline development defect at 25 °C that produces germline-ablated adults ^18,19^. All strains were maintained on NGM agar plates seeded with *E. coli* NEC937 B (OP50 ΔuvrA; KanR) ^20^.

### Animal handling

Animals were cultured in standard conditions ^21^. Wildtype and fertile *glp-1(e2141*) animals were cultivated at 20 °C, and synchronized at L4 via lay-off. Sterile *glp-1(e2141*) animals were developed and cultivated at 25 °C, and equally synchronized at L4 via lay-off. Lysis to obtain mRNA was performed using the protocol described by Serra *et al*. ^22^ with the addition of ERCC spike-ins to the lysis buffer to reach a final dilution of 1:40000. For bulk samples, 15 animals were placed together into 120 µL of lysis buffer. For single-individual experiments, 1 animal was placed into 8 µL of lysis buffer for individual transcriptomics.

### Multi-generational gene-expression protocol

To systematically identify heritable gene-expression states, we collected the transcriptomes of 96 siblings on the second day of adulthood while taking a subset of 24 for multi-generational analysis. For each of these 24 individuals, immediately before sample collection, we collected every progeny laid in a four-hour period. On average, 43 eggs were laid per parent, leading to a total population of 1045. We let these cousins age until the 2^nd^ day of adulthood and sacrificed them for transcriptomics at the same age as we had sacrificed their parents.

### Comparing germline-ablated vs wild-type populations

For Smart-seq2-based STOMP-seq, single-individual samples were obtained for 48 wild-type and 48 *glp-1(e2141*) individuals. After lysis, each individuals’ lysate was split into two aliquots—one processed with Smart-seq2 and the other using STOMP-seq. In this case, STOMP-seq was performed on samples organized into eight separate pools, each containing 6 wild-type and 6 *glp-1(e2141*) individuals (Fig. 4a). For Smart-seq3-based STOMP-seq, we collected 3 population pools of wild-type and *glp-1(e2141*) individuals each. After lysis, each pools’ lysate was split into aliquots—one processed with Smart-seq3 and the other using 12 different STOMP-seq barcoded TSOs. In this case, STOMP-seq was performed on samples organized into six separate pools, each containing 6 wild-type and 6 *glp-1(e2141*) population pools (Fig. 4a). This experimental design allows for the detection of a variety of potential technical artifacts introduced by the sequencing protocol—including barcode switching and barcode introduced biases—which necessarily degrade the protocol’s ability to identify gene-expression differences between wild-type and *glp-1(e2141*) individuals.

### Novel Programs, Software, Algorithms

Data analysis code is made available in the github repository https://github.com/nstroustrup/AID-engineering

### Web Sites/Data Base Referencing

The *C. elegans* reference genome (release WS265) was obtained from Wormbase (www.wormbase.org).

## Results

### 5’-end sample barcoding enables early multiplexing

Following a 5’ barcoding strategy (Fig. 1a,b; S1a,b), we can implement an RNA-seq protocol that multiplexes samples immediately after RT, reducing liquid-handling 12-fold while increasing transcriptome coverage compared to Smart-seq2. By multiplexing twelve samples into a single well immediately after RT, we lower twelve-fold the effective sample size during liquid handling, which reduces “hands-on” time actively handling liquids from 2.9 hours using Smart-seq2 down to 30 minutes per plate using STOMP-seq (Table S1). For an experiment involving twelve or more 96-well plates, our approach further reduces “hands-off” time spent waiting for PCRs and incubations to complete from 7.8 hours to 3.3 hours per plate. In total, our approach increases the overall protocol throughput more than 4-fold (Fig. 1c). We also reduce the number of Nextera indexing primers needed to barcode samples, which can be a major expense for newer, large-capacity sequencing machines. By making use of these flow cells more practical, our 5’ barcoding strategy further reduces the per-base sequencing costs by two-fold or more.

**Figure 1.**
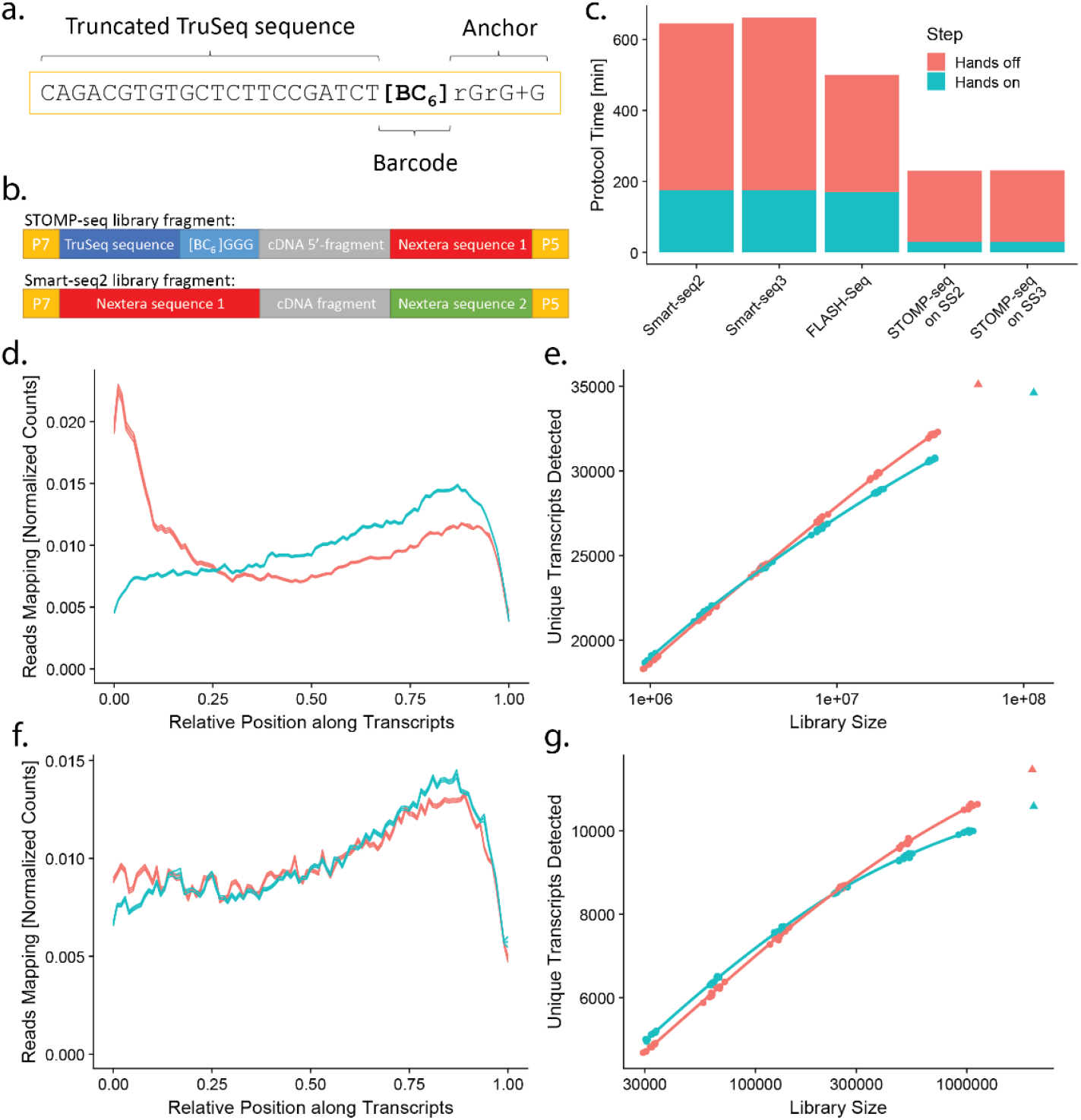
The STOMP-seq protocol enables a fast early-multiplexing strategy. **a**. Composition of the STOMP-seq Template Switching Oligo (TSO) with TruSeq sequence and barcode. **b**. In comparison to Smart-seq2 (*bottom*), STOMP-seq (*top*) generates 5’-end barcoded cDNA, enabling a multiplexing strategy that **c**. dramatically decreases active (*blue*) and passive (*red*) protocol execution times. **d**. Average read mapping position along the thousand most abundant transcripts in STOMP-seq (red) and Smart-seq2 (*blue*) libraries. **e**. The number of unique transcripts detected in STOMP-seq (*red*) and Smart-seq2 (blue) libraries of different sizes. **f**. Average read mapping position along the thousand most abundant transcripts in STOMP-seq (*red*) and Smart-seq3 (blue) libraries. **g**. The number of unique transcripts detected in STOMP-seq (*red*) and Smart-seq3 (*blue*) libraries of different sizes.

To evaluate our approach, we collected mRNA from individual *C. elegans* using either STOMP-seq or Smart-seq2. First, we looked at the fragment size distribution of libraries produced using either method, and identified a peak that corresponded to the disproportionate amplification of mRNA derived from genomic regions whose sequences showed similarity to the TruSeq “read2” primer sequence (Fig. S1c). In fact, 56.5% of 5’-barcoded reads and 5.7% of Smart-seq2 reads contained sequences within a 3-basepair mismatch of the TruSeq primer sequence fragment “CACGTCTGAAC”. These results suggested that this TruSeq primer fragment favorably recruits Tn5 enzymes during tagmentation, resulting in the formation of short, sense-less, or even partly un-sequencable library fragments by tagmenting inside the “read2” sequencing initiation sequence. Consequentially, 22.7% of paired-end 5’-barcoded reads completely lacked “read2”, a phenomenon exacerbated by our strategy over Smart-seq2 due to our inclusion of this Tn5 bias sequence on 5’ TSOs.

To eliminate this technical bias and stop the TruSeq primers from selectively recruiting Tn5 enzymes during tagmentation, we altered our TSO sequence to include only the last 22 bp of the full 34 bp TruSeq sequence (Fig. 1a, S1c, Table S4). This reduced the prevalence of TruSeq primer complementary reads to 14.3% and reduced paired end reads lacking “read2” to 8.1%. As the full TruSeq sequence is required later during sequencing using Illumina flow cells, we add the full sequence back during the subsequent library amplification step using modified N7 indexing primers (Table S4).

### Early pooling maintains library coverage and replicability

We first assessed the quality of cDNA libraries produced by Smart-seq2-based STOMP-seq. In contrast to the 3’ bias in read coverage often observed in Smart-seq2 ^23^, STOMP-seq produces full-length cDNAs with a moderate bias toward the 5’-end (Fig. 1d). At equivalent read depths, STOMP-seq libraries contain a higher number of unique transcripts compared to Smart-seq2 (Fig. 1e), a beneficial result of STOMP-seq amplifying only the single 5’ fragment of each mRNA. We then assessed the quality of cDNA libraries produced by Smart-seq3-based STOMP-seq, finding that both protocols produce a similar full length coverage of transcripts, with STOMP-seq showing reduced drop-off in 5’-end coverage compared to Smart-seq3 (Fig. 1f). Similar to the effects observed on the Smart-seq2 protocol, Smart-seq3-based STOMP-seq libraries contain a higher number of unique transcripts compared to the original Smart-seq3 (Fig. 1g).

Technical replicates analyzed with STOMP-seq and Smart-seq2 show equivalently high inter-replicate correlations, 98.8% and 97.9% (Fig. S1d) respectively, showing that our modified TSOs do not alter the accuracy of sequencing results. We observed a somewhat lower correlation between the two protocols performed on the same samples (90.2%, Fig. S1d), which is entirely explained by the transcript length bias present in Smart-seq2 (Fig. S1e) but eliminated by STOMP-seq’s 5’ sequencing strategy. In comparison to Smart-seq2, we see a lower inter-replicate correlation of Smart-seq3-based STOMP-seq (92.7%, Fig S1f), while the correlation between the two protocols performed on the same samples is with 90.6% comparable to that between STOMP-seq and Smart-seq2.

### Selecting a set of 5’ barcodes to ensure homogeneous amplification across samples

To identify any bias introduced by specific Smart-seq2-based STOMP-seq barcode sequences into amplified cDNA libraries, we processed three replicate aliquots taken from a single wild-type biological sample using each of 21 TSO barcode sequences processed in separate (un-pooled) reactions. We identify a set of 12 TSO barcodes that produce cDNA yields 0.76 ± 0.2 of that obtained using Smart-seq2 (Fig. S1g), generating balanced multiplexed cDNA libraries (Fig. S1h). Across all these barcodes, at least 89% of sample reads contained identifiable 5’ barcodes (Fig. S1i).

We then performed principal component (PC) analysis to identify any influence of TSO barcodes on cDNA library composition. As would be expected from 18*3 technical replicates measured from the same sample, most expression variance between technical replicates took the form of unstructured noise (Fig. S2a, Supplementary Note 2). However, we identified two PCs attributable to the influence of TSO barcode biases (Fig. 2a): PC1 (5.38%) is produced by differential rates of TSO “mis-priming”, in which barcode sequences modulate the rate of an “off-target” binding of TSOs to mRNA strands to initiate reverse transcription and generate aberrant non-terminal PCR amplicons ^19^ (Fig. 2c diagram; Supplementary Note 1). PC2 (4.2%) is produced by “strand invasion” of TSOs, which occurs when the RT enzyme switches templates from mRNA to the TSO prematurely before reaching that mRNA’s 5’-end ^24^ (Fig. 2c diagram). We considered the possibility that the rates of TSO strand invasion might be modulated by the sequence of TSO barcodes, leading to some TSOs exhibiting higher rates of strand invasion than others. If such a phenomenon were present, then barcodes with higher rates of strand invasion would generate higher proportions of truncated transcripts, which, after tagmentation, produce reads that map closer to the 3’-end of mRNAs. Looking for a bias in read mapping positions along transcripts, we found that differential rates of strand invasion between barcodes produce differences in the average read mapping position between the 3’ and 5’ ends of a transcript’s mapping position (Fig. S2f). These differences correlate (rho_S_ = .61) to the second PC of our sequenced libraries (Fig. 2i), demonstrating that differential rates of strand invasion drive 4.2% of all inter-sample variance generated by barcodes. We obtained similar results when analyzing Smart-seq3-based STOMP-seq. Selecting the 12 best-performing Smart-seq2-based STOMP-seq barcodes, we prepared triplicate samples using Smart-seq3-based STOMP-seq. We find that different barcode sequences separate samples along two PCs (Fig. S3a): PC1 (18.3%) and PC3 (6.83%). The effect separating the barcodes along PC1 (Fig. S3b) results from both TSO “mis-priming” and “strand invasion” phenomena we saw in Smart-seq2-based STOMP-seq (Fig 2k,l, S3c).

**Figure 2.**
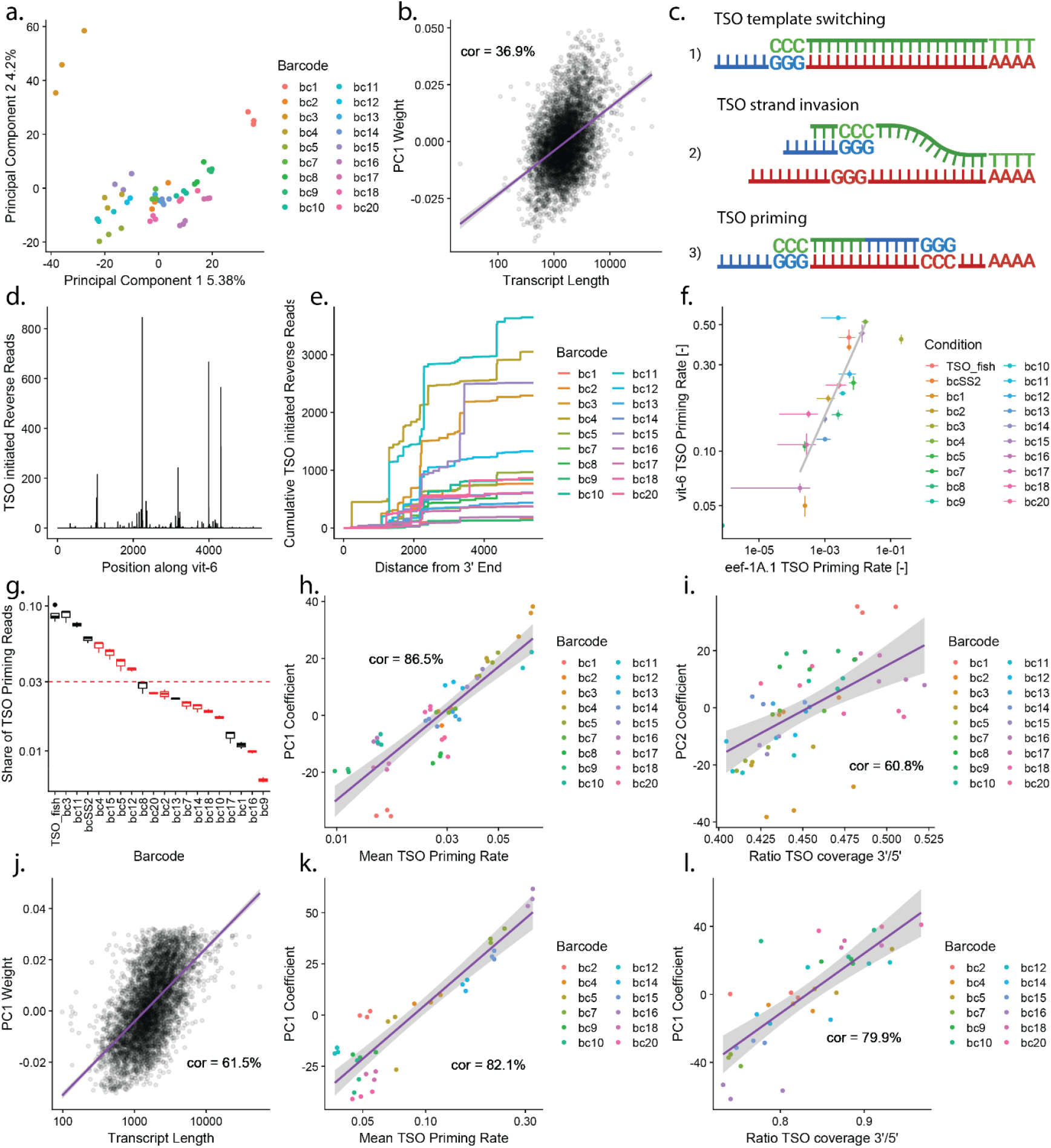
The effects of TSO barcode sequence on STOMP-seq cDNA libraries. **a**. Three replicates were performed using each of 18 different TSO barcode sequences on identical aliquots taken from the same biological sample. Principal component analysis identifies two main axes of technical noise contributing 5.4% and 2.4% of all inter-sample variation, respectively. **b**. Transcript PC1 weights plotted against the length of the respective transcripts, with linear model fit (*purple*). **c**. Scheme of potential TSO interactions with the mRNA. 1) Completion of the first strand cDNA through template switching, 2) strand invasion, and 3) priming of the reverse transcription. **d**. The coverage of reads from cDNA sequences adjacent to the Smart-seq2 TSO, resulting from aberrant TSO priming along vit-6. **e**. The cumulative number of aberrant TSO priming reads produced by each barcode (*colors*) along vit-6. **f**. The fraction of reads derived from aberrant TSO priming compared for two of the most highly-expressed genes, eef-1A.1 and vit-6, measured across all TSO barcodes in triplicate technical replicates, with standard error bars. **g**. The global share of reads generated by TSO priming, separated by triplicates of each barcode and with the mean share across the barcodes (*red dotted line*). Two strategies, TSO_fish and bcSS2, try to assess the distribution of TSO priming among transcripts processed with Smart-seq2. The final 12 chosen STOMP-seq barcode sequences are highlighted in red. **h**. The position of each sample on PC1, i.e. the PC1 coefficients, are compared to the average rate of TSO priming for a biological sample when amplified with different TSO barcode sequences (*colors*), with linear fit (*purple*). **i**. The ratio of 3’ to 5’-end coverage of forward reads plotted against PC2 coefficients for barcoded samples in triplicates, with linear model fit (purple). **j**. Transcript PC1 weights plotted against the length of the respective transcripts, with linear model fit (*purple*). k. The position of each sample on PC1, i.e. the PC1 coefficients, are compared to the average rate of TSO priming for a biological sample when amplified with Smart-seq3-based STOMP-seq using different TSO barcode sequences (*colors*), with linear fit (*purple*). **l**. The ratio of 3’ to 5’-end coverage of forward reads plotted against PC2 coefficients for barcoded Smart-seq3-based STOMP-seq samples in triplicates, with linear model fit (purple).

In both Smart-seq2- and Smart-seq3-based STOMP-seq, the influence of such small but measurable biases can be effectively eliminated by adopting a barcode permutation strategy. Any confounding effect of barcode sequences can be eliminated by ensuring that barcode sequences do not co-vary with sample type: ensuring that technical replicates of each covariate group are spread across different barcode sequences. Without barcode permutation, barcode biases substantially inflated the false-positive rate (FPR) of tests for statistically-significant changes in any gene’s expression between technical replicates of identical samples (Fig. S2g). After barcode permutation, the FPR decreased to 10^-3^ % on average (Fig. 3a,b), demonstrating an absence of barcode bias after permutation. This permutation strategy is equally effective in dropping FPR when applying STOMP-seq on Smart-seq3 (Fig 3c,d, S3d).

**Figure 3.**
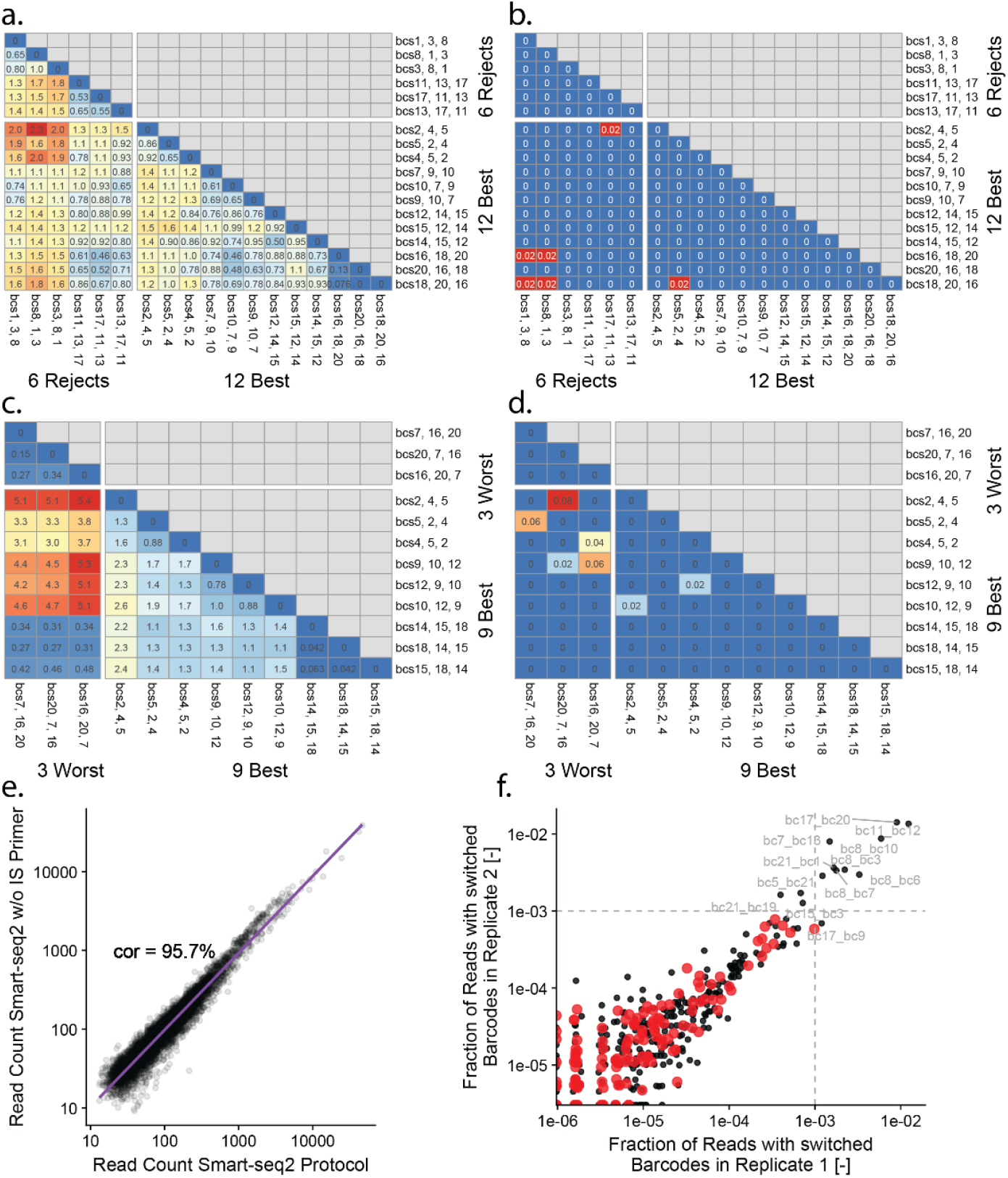
Barcode effects can be minimized by barcode selection and combination. **a**. Heatmap of false positive differentially expressed genes inflation over false positive rates expected for p<.01, identified using triplicates of identical samples amplified using combinations of three different barcodes. Comparisons between the final 12 chosen STOMP-seq barcode sequences are grouped together on the bottom right. **b**. The false positive (FP) rates in differential expression analysis (FP per 10,000 transcripts at p.adjusted<.01) of identical samples processed with different STOMP-seq barcodes. Comparisons between the final 12 chosen STOMP-seq barcode sequences are grouped together on the bottom right. **c**. Heatmap of false positive differentially expressed genes inflation over false positive rates expected for p<.01, identified using triplicates of identical samples amplified with Smart-seq3-based STOMP-seq and combinations of three different barcodes. Comparisons between the 9 best-performing STOMP-seq barcode sequences are grouped together on the bottom right. **d**. The false positive (FP) rates in differential expression analysis (FP per 10,000 transcripts at p.adjusted<.01) of identical samples processed with Smart-seq3-based STOMP-seq and different STOMP-seq barcodes. Comparisons between the 9 best-performing STOMP-seq barcode sequences are grouped together on the bottom right. **e**. Correlation of cDNA profiles generated with or without the addition of IS primers for first-strand cDNA amplification, with linear model fit (*purple*). **f**. The rates of barcode swapping between STOMP-seq barcode pairs are compared across two replicates, with the best 12 barcodes highlighted (*red*).

**Figure 4.**
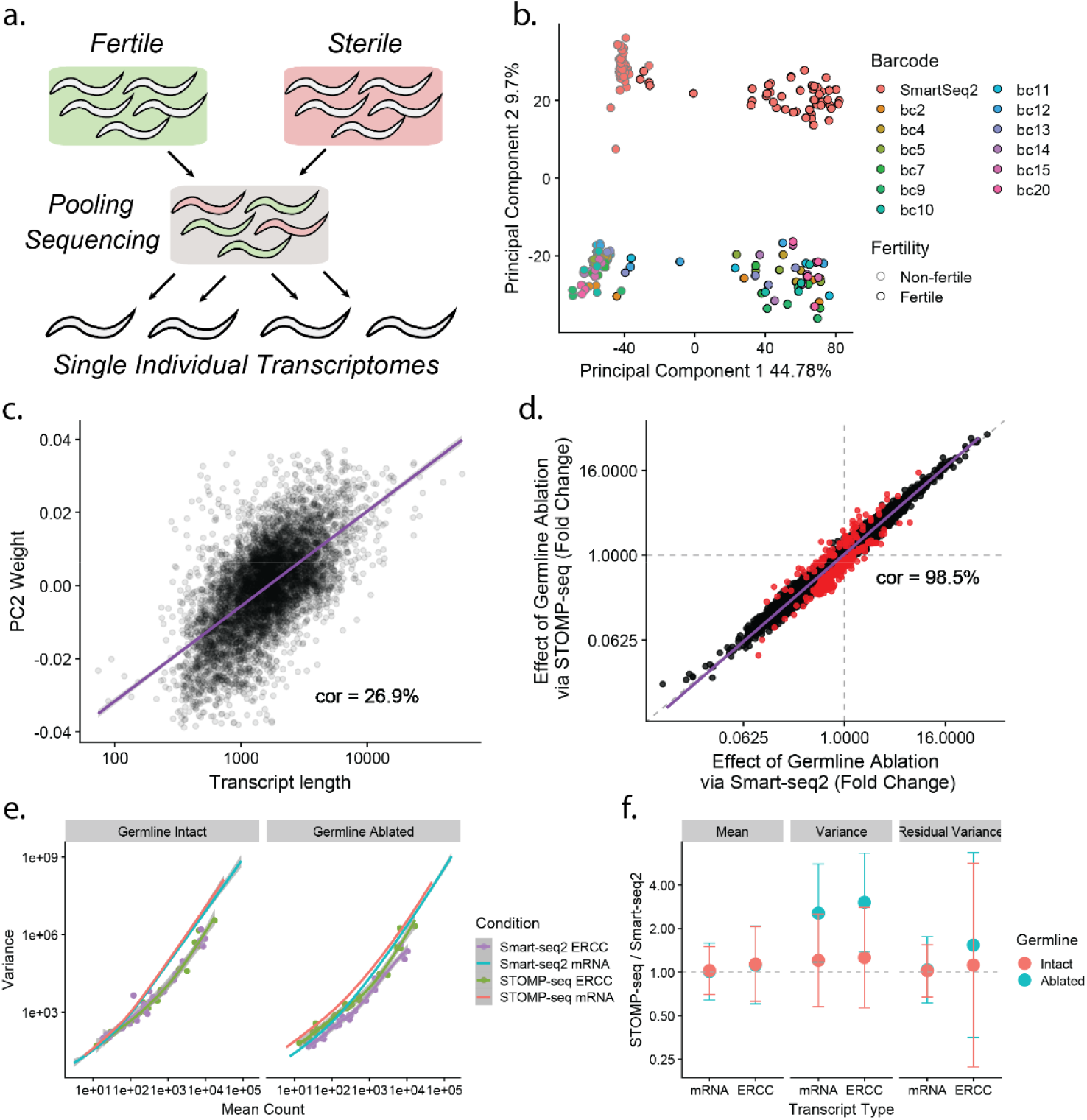
STOMP-seq accurately quantifies biological differences between samples. **a**. Experimental layout of the early pooling assessment with fertile and sterile glp-1(e2141) individuals, using barcoded samples. **b**. PCA of the transcriptomes of 48 intact (*black outline*) and 48 germline-ablated *glp-1(e2141) (no outline)* individuals, processed either in STOMP-seq pools (*colors*) or using Smart-seq2 (*red*). **c**. The PC2 weight of each transcript plotted against its length, with linear model fit (*purple*). **d**. The fold-change effect of germline ablation on each gene’s expression, measured by Smart-seq2 (x-axis) and STOMP-seq (y axis), with significant differences between protocols at p<.01 highlighted red, and with linear fit (*purple*). **e**. The variance in transcript counts plotted against the transcript’s mean count for fertile (*left*) and sterile (*right*) animal samples. Samples are separated by protocol and by technical and biological variance. For visibility, only spline fits are displayed for biological variance. **f**. Summary statistics for the relative abundance of transcripts (*Mean, left*), the inter-sample variance of transcripts (*Variance, center*), and the residual variance (*Residual Variance, right*), estimated using parametric methods as described in Eder *et al*. 2024. Statistics shown are the average across all mRNAs and ERCC transcripts measured in intact (*red*) and germline-ablated populations (*blue*), with error bars indicating one standard deviation.

**Figure 5.**
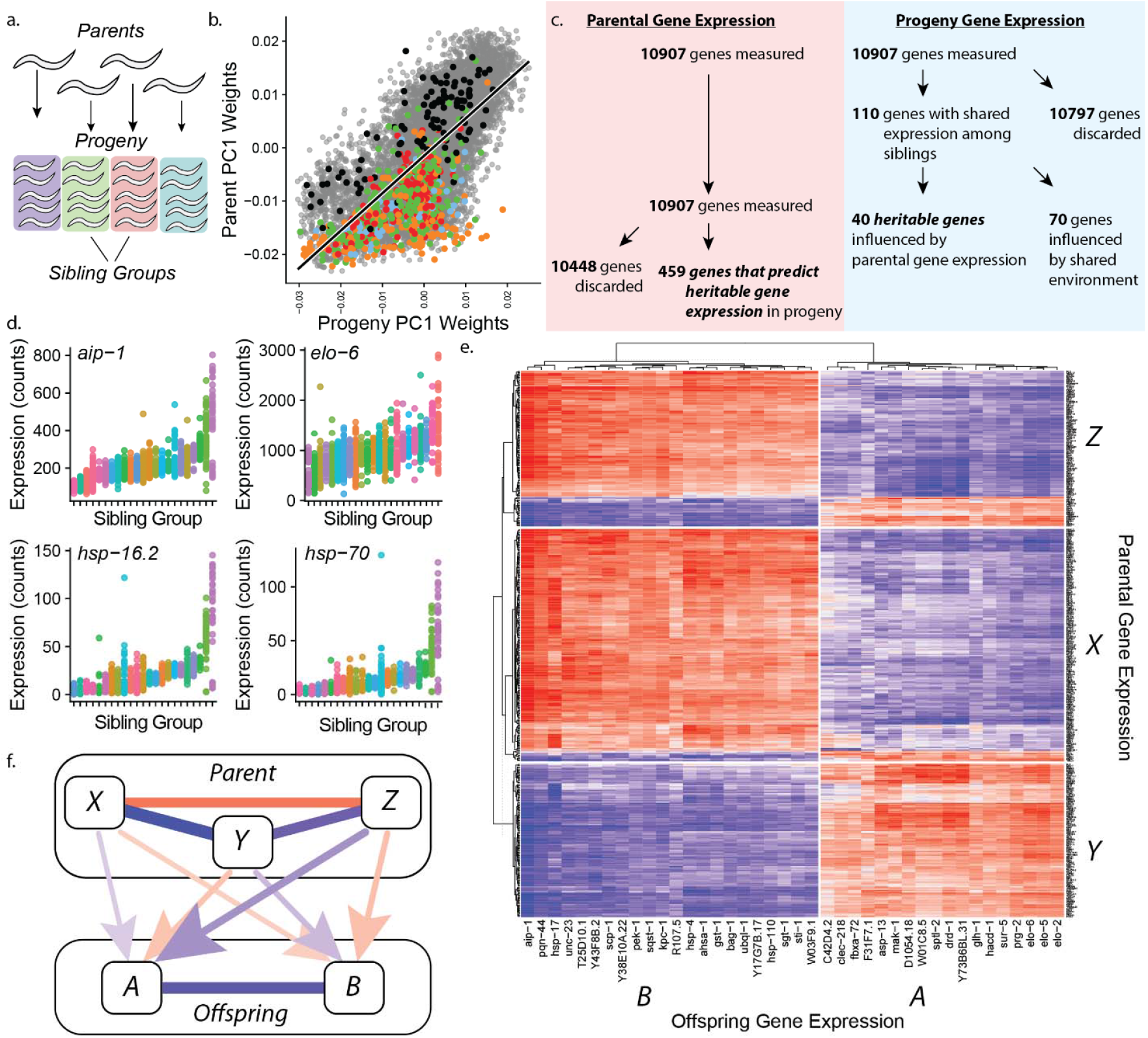
Using population-scale STOMP-seq to identify heritable gene-expression states. **a**. Transcriptomes of 24 parents and their 1045 progeny were collected. **b**. Exhibited correlated first principal components, with germline-expressed genes (*black*) anti-correlated with somatically-expressed genes (*colors*). **c**. Workflow for detecting heritable gene expression through linear regression and partial least squares regression analysis. **d**. Four example genes whose expression is more similar between siblings (*colors*) than cousins. **e**. Hierarchical clustering of heritable genes reveals two gene groups (A, B) whose expression in progeny associates with the expression of three different groups (X,Y,Z) in their parents. Colors indicate Pearson correlation between −1 (*blue*) and 1 (*red*). **f**. The full causal model of gene-expression heritability, involving three sets of parental genes driving adult gene-expression of two groups of genes in offspring, with line color and width proportional to the correlation strength.

### Selecting a set of 5’ barcodes to minimize index hopping

We then considered the potential for “barcode-switching”, also known as “index hopping”, in which transcripts from one sample inadvertently acquire another sample’s barcode during multiplexed PCR amplification. We demonstrated the potential for “barcode-switching” by performing an experiment in which PCR amplification after RT was performed without adding the standard cDNA amplification primers. Without these primers, the PCR reactions nevertheless produced usable cDNA libraries (Fig. 3e), highlighting the potential for 5’ TSO primers remaining from RT to act as primers during PCR, based on their complementary TSO sequence already appended to each cDNA. This reaction is no problem when barcode sequences of the 5’ TSOs acting as primers are a perfect sequence match for the TSO sequence of the cDNA. However, it becomes problematic if barcoded TSOs with partial complementarity to the cDNA act as primers, effectively swapping the correct barcode on the cDNA that was added during RT for an incorrect barcode present during PCR amplification.

To estimate the worst-case rates of actual barcode switching within the STOMP-seq protocol, we performed first strand cDNA amplification of transcripts barcoded with each barcode during RT in the presence of a surplus of 5’ TSOs containing each of the other barcode sequences. Each combination of cDNA barcode and 5’ TSO barcode mix was performed in a separate well, allowing us to identify barcode switching computationally in the resulting sequencing libraries as transcripts barcoded by another 5’ TSO’s sequence rather than the original cDNA barcode. In this way, we identify a set of 12 barcode sequences with switching rates all lower than 0.078% (Fig. 3f). We conclude that the true STOMP-seq barcode-switching rate is lower than the inherent error rate of the underlying illumina sequencing technology ^25^, and therefore cannot be lowered further.

### Evaluating STOMP-seq for differential expression analysis

To demonstrate the performance of Smart-seq2-based STOMP-seq in differential expression analysis, we sequenced a population of 48 wild-type and 48 germline-ablated ^18^ individuals using STOMP-seq in 12x multiplexed mode as well as standard Smart-seq2 (Fig. 4a). We observe the same, clear separation of wild-type and germline-ablated *glp-1(e2141*) individuals along PC1, explaining 44.78% of all sample variance (Fig. 4b). PC2 reflects the inherent length bias of Smart-seq2 relative to STOMP-seq (Fig. 4c). The two methods produced extremely similar estimates of gene-expression changes between wild-type and *glp-1(e2141*) individuals (rho_S_ = .99) (Fig. 4d). Only 328 genes, 5.09% of the transcriptome, showed statistically significant but very small magnitude differences in effect sizes measured by each method. We then considered Smart-seq3-based STOMP-seq in a similar 12x multiplexed differential expression analysis. We again observe a clear separation of wild-type and germline-ablated *glp-1(e2141*) individuals along PC1, explaining 33.01% of all sample variance (Fig S3e). STOMP-seq produced very similar estimates to Smart-seq3 for the magnitude of gene-expression changes between wild-type and *glp-1(e2141*) individuals (rho_S_ = .90), and no genes showed statistically significant differences in effect sizes measured by each method (Fig. S3f). Together, these data demonstrate the suitability of STOMP-seq as a “drop-in” replacement for both Smart-seq2 and Smart-seq3 protocols.

### Applying STOMP-seq for population-scale, multi-generational, single-individual sequencing

Taking advantage of the four-fold increase in sample processing speed of STOMP-seq, we then performed a multi-generational study of gene expression in *C. elegans* designed to identify heritable gene-expression states passed down from parent to progeny. In *C. elegans*, transgenerational inherence allows parents to modulate a wide range of phenotypes in their genetically-identical progeny ^26–28^, but the potential for inheritance of gene-expression states in the absence of genetic variation remains unexplored due to the large population sizes required to characterize such phenomena.

To systematically identify heritable gene-expression states, we used STOMP-seq to perform a population-scale RNA-sequencing experiment, collecting 1141 individual transcriptomes that measure gene-expression differences across a large family tree including 96 siblings from a parental generation and on average 43 of the adult progeny (cousins) from each of 24 of the parents (Fig. 5a). To identify gene-expression changes that persist for one full generation, we measured gene-expression of the progeny during their adulthood, at the exact same age at which gene-expression had been measured in their parents. We find that inter-individual gene-expression variability takes a similar form across the two generations, with the first PC weights correlating at rho = .71 (Fig. 5b, Fig. S4a), demonstrating a varying ratio of somatic to germline mRNA previously shown as the largest source of physiologic heterogeneity in this species ^19^. The remaining PCs showed varying degrees of correlation between the two generations (Fig. S4a-b), which we interpret as suggesting that co-variation among similar groups of genes contribute un-equally to inter-individual variation across the two generations.

To identify aspects of gene-expression variance resulting from heritable factors, we first searched for genes whose expression was more similar between siblings than between cousins. Using categorical regression, we identified 110 such genes, or about 1% of all genes measured in the experiment (Fig 5c,d; Fig. S4c,d). Remarkably, this gene set contained both *hsp-16*.*2* and *elo-6*, which are members of a small group of genes whose protein abundance predicts remaining lifespan within isogenic populations of *C. elegans* ^29,30^—suggesting that these genes ability to act as aging biomarkers derive in part from their transduction of heritable factors.

The gene-expression similarities among siblings that we identify could be the result of inherited factors, but they also might result from environmental factors shared among siblings ^31^. Therefore, although all individuals were housed on identically-prepared culturing plates, any residual environmental differences might act as confounding factors in our heritability analysis. To address this problem, we reasoned that it should be possible to distinguish inherited gene-expression states from environmental effects by relating progeny gene-expression states to parental gene expression states. Since parents and progeny occupy the same environment only for a four-hour window during the parents’ adulthood and the progeny’s late embryonic development, we reasoned that similarities among siblings that showed quantitative correlations with their parent’s gene-expression states would be more likely to represent inherited factors whose transmission is not mediated by environmental factors.

To identify such factors, we performed partial least squares regression and identified 40 genes out of the original 110 whose shared expression in sibling groups was significantly associated with quantitative changes in gene expression in their parents (Fig. 5c,e, S4e,h). For the remaining 70 genes lacking strong intergenerational correlations, we conclude that either environmental factors shared among sibling groups or aspects of parental physiology not measurable using our transcriptomic method are responsible for gene-expression similarities among siblings.

These 40 genes with inter-generational correlations were partially but not completely correlated with each other across progeny, grouping into at least three distinct co-varying, partially coupled sets (Fig. 5e,f; Fig. S4e,f). We interpret these results as suggesting that three partially coupled aspects of physiology in parents transmit a heritable signal that modulates the adult gene-expression of their progeny. Remarkably, these genes’ expressions do not appear to be directly heritable—e.g., expression of any particular gene in the parent does not predict the expression of that same gene in progeny (Fig. S4i). Rather, heritability appears to act in *trans*, in which three partially-coupled sets of genes in parents, X,Y,Z influence two distinct, non-overlapping sets of genes in the progeny, A and B (Fig. 5f).

Therefore, we can assemble our data into an inter-generational map of heritable gene-expression variance: gene sets X,Y,Z that are correlated with two sets of co-varying genes in the progeny – A and B. At the group level, X,Y, and Z are highly but incompletely correlated (rho(X,Y) = −.94, rho(X,Z) = .68, rho(Y,Z) = −.74; Fig. 5c), as are A and B (rho(A,B) = −.83; Fig. 5b). The inter-generational correlations are lower than the within-generational correlations (rho(X,A) = .23, rho(X,B) = .24, rho(Y,A) = .3, rho(Y,B)= −.29, rho(Z,A) = −.48, rho(Z,B) = .32; Fig. 5a), as might be expected because the causal relationship between generations must be established during early embryogenesis of the progeny and then persist until adulthood.

Our results comprehensively reveal the ways in which intrinsic sources of organismal variation in gene-expression are propagated across generations, potentially leading to inter-generational couplings in phenotypic variation even absent any genetic variation.

In conclusion, STOMP-seq enables high-throughput, flexible plate-based experimental designs, amenable for manual or robot protocols—a “drop-in” multiplexed replacement for SMART RNA-seq protocols that reduces protocol time while improving quality.

## Discussion

Here, we present STOMP-seq, an RNA-sequencing protocol that increases the throughput of the SMART sequencing protocols Smart-seq2 and Smart-seq3 while preserving their precision and sensitivity. Central to our approach is a sample-identifying barcode located on the Template Switching Oligo (TSO) used to introduce this barcode to the 5’-end of each transcript. By systematically identifying and eliminating the specific molecular mechanisms reducing the fidelity of reverse transcription, PCR amplification, and tagmentation, we produce a highly-multiplexed protocol with performance equivalent to or slightly exceeding that of Smart-seq2 and Smart-seq3. STOMP-seq reduces hand-on time during sample preparation by up to 75% compared to the published SMART protocols while retaining equivalent or better full-length coverage, replicability, and accuracy.

We identify and solve an unfortunate molecular mechanism present in the original Smart-seq2 protocol: TSOs substitute for poly-dT primers during reverse transcription, resulting in 8% of total reads and reads from 89% of all unique transcripts mapping to truncated transcripts—a substantial deviation from the full-length reads promised by the protocol. This molecular mechanism should be consistent across all samples processed by Smart-seq2 and will therefore not act as a confounding factor in analyses performed exclusively on Smart-seq2 data and no other sequencing technologies. However, the mechanism represents a fundamental challenge to any protocol development that involves change to the 5’ TSO, which modulate the mechanism and turn it into a serious confounding factor. Our protocol suppresses the TSO priming mechanism—a crucial step in the development of STOMP-seq.

STOMP-seq opens the door to highly-multiplexed plate-based experimental designs. We demonstrate the potential here in a multi-generational, population-scale RNA-sequencing study, which identifies a set of genes whose expression in adults appears to be partially determined by measurable physiologic states in the respective parent.

STOMP-seq is amenable to use by hand and by robot, minimizing liquid handling in any context. Importantly, STOMP-seq allows users to multiplex multiple samples across single sequencing barcodes allowing a larger number of samples to be loaded in a single flow cell. Pooling more samples per flow cell in turn allows users to move to larger-capacity flow cells with lower per-read costs. In summary, STOMP-seq is a drop-in replacement for RNA-seq protocols of the SMART family that dramatically reduces the processing time and cost of high-quality, plate-based RNA-sequencing.

## Data and Code Availability

All sequencing data will be made available through the NIH Sequence Read Archive (SRA). All data analytic code is available in the repository https://github.com/nstroustrup/STOMP-seq.

## Author Contributions

ME led the investigation and data curation; NS led the supervision and software development, and contributed to data curation; Both authors contributed equally to conceptualization, methodology, analysis, and writing of the manuscript.

## Acknowledgements

We thank Jochen Hecht and all members of the Dynamics of Living Systems group for discussions and encouragement throughout this project. We thank Joy Alcedo (Wayne State University) for nematode strains. Some strains were provided by the CGC, which is funded by NIH Office of Research Infrastructure Programs (P40 OD010440).

## Supplementary Information

### Supplementary Notes

#### Supplementary Note 1 – Internal TSO priming

To understand the influence of 5’ TSO barcodes on library composition, we considered the barcode effects captured by inter-sample variance along PC1 (Fig. 2a, S2a, S3a,b). The weights assigned to genes for PC1 are negatively correlated with the length of those genes’ coding sequences (rho_S_ = −.37 for Smart-seq2-based STOMP-seq and rho_S_ = −.62 for Smart-seq3-based STOMP-seq respectively; Fig. 2b,j), suggesting that barcodes vary in the extent to which they favor shorter transcripts over longer transcripts. Furthermore, the barcodes that favor longer transcripts (low PC1 values) also tend to produce overall higher cDNA yields (Fig. S2b), suggesting that these barcodes generate more cDNA overall in proportion to gene length. Such a phenomenon is reminiscent of the recently described “aberrant TSO priming” activity present in the Smart-seq2 protocol: previously we had described that Smart-seq2 TSOs can bind to non-terminal CCC sites within the coding sequence of specific mRNAs, catalyzing the generation of extra transcripts beyond what would normally be generated by the TSO’s expected template-switching activity ^1^. Since this TSO priming activity is observed in the standard Smart-seq2 TSO, we hypothesized that our incorporation of TSO barcodes might modulate this mechanism, with different barcodes producing different rates of internal TSO priming. Such differing rates of internal TSO priming would then result in some barcodes favoring longer transcripts, as the probability of internal TSO priming will increase proportionally to transcript length—the probability of a transcript containing a sequence complementary to the TSO barcode will increase in proportion to that transcript’s length.

A technical feature of STOMP-seq allows us to systematically measure internal TSO priming rates and test this hypothesis. Since sequencing of every transcript’s “read2” is initiated by the TSO, we can detect reads derived from internal TSO priming based on their unique signature: a TSO sequence incorporated anti-sense relative to the transcript’s cDNA sequence (Fig. 2c diagram). We first considered a highly-expressed gene, *vit-6*, and found peaks in read coverage distributed across the transcript (Fig. S2c). These peaks were formed almost exclusively from reads derived from internal TSO priming (Fig. 2d) rather than cDNA completion by the TSO (Fig. S2d). The rates of internal priming at these peaks vary across barcodes (Fig. 2e): The extend of internal priming at any given position varies with the similarity of the barcode sequences to the transcript’s sequence downstream of a potential CCC priming site (Fig. S2e, Tab. S2). Furthermore, the total amount of internal priming across the barcodes also varies between transcripts (Fig. 2f). On average across all barcodes, TSO internal priming is responsible for roughly 3.53% of all reads in our libraries, lower than the 8.71% we find are produced by the Smart-seq2 protocol (Fig. 2g). Across barcodes, the fraction of reads derived from internal TSO priming is strongly correlated (rho_S_ = −.87) to the first PC of our sequenced libraries (Fig. 2h), demonstrating that differential rates of internal TSO priming generate 5.38% of inter-sample variance—the single largest influence of individual barcode sequences to library composition.

In summary, we find that variation along the first PC is a reaction present in Smart-seq2 in which TSOs can bind to non-terminal CCC sites within the coding sequence of specific mRNAs, catalyzing the generation of extra transcripts beyond what would normally be generated by the TSO’s expected template-switching activity. On average across all barcodes, we find that TSO internal priming is responsible for roughly 3.53% of all reads in our libraries, substantially lower than the 8.71% of transcripts produced by TSO internal priming in the Smart-seq2 protocol.

#### Supplementary Note 2 – Quantifying Inter-sample Variance

Inter-sample variance, introduced as technical noise by a sequencing method, is important to applications seeking to measure biological variation between samples, for example in aging studies ^1^. We therefore estimated the inter-sample variance of each gene across our set of identical aliquots processed with either STOMP-seq or Smart-seq2. Inter-sample variance depends on the mean expression of each gene, so we estimated biases between the two methods in respect to gene-expression mean, gene-expression variance, and residual variance that uses ERCC spike-ins to remove technical noise from variance estimates. We saw no bias in gene-expression mean between STOMP-seq and Smart-seq2 (Fig. 4e,f). STOMP-seq showed a slightly elevated inter-sample variance in one out of two conditions, but this occurred for both ERCC and mRNA derived transcripts, and therefore we saw no difference in residual variance between STOMP-seq and Smart-seq2 across either condition. We therefore conclude that STOMP-seq is equivalent to Smart-seq2 in its ability to measure true biological variation between samples.

### Supplementary Tables

**Supplementary Table 1.**
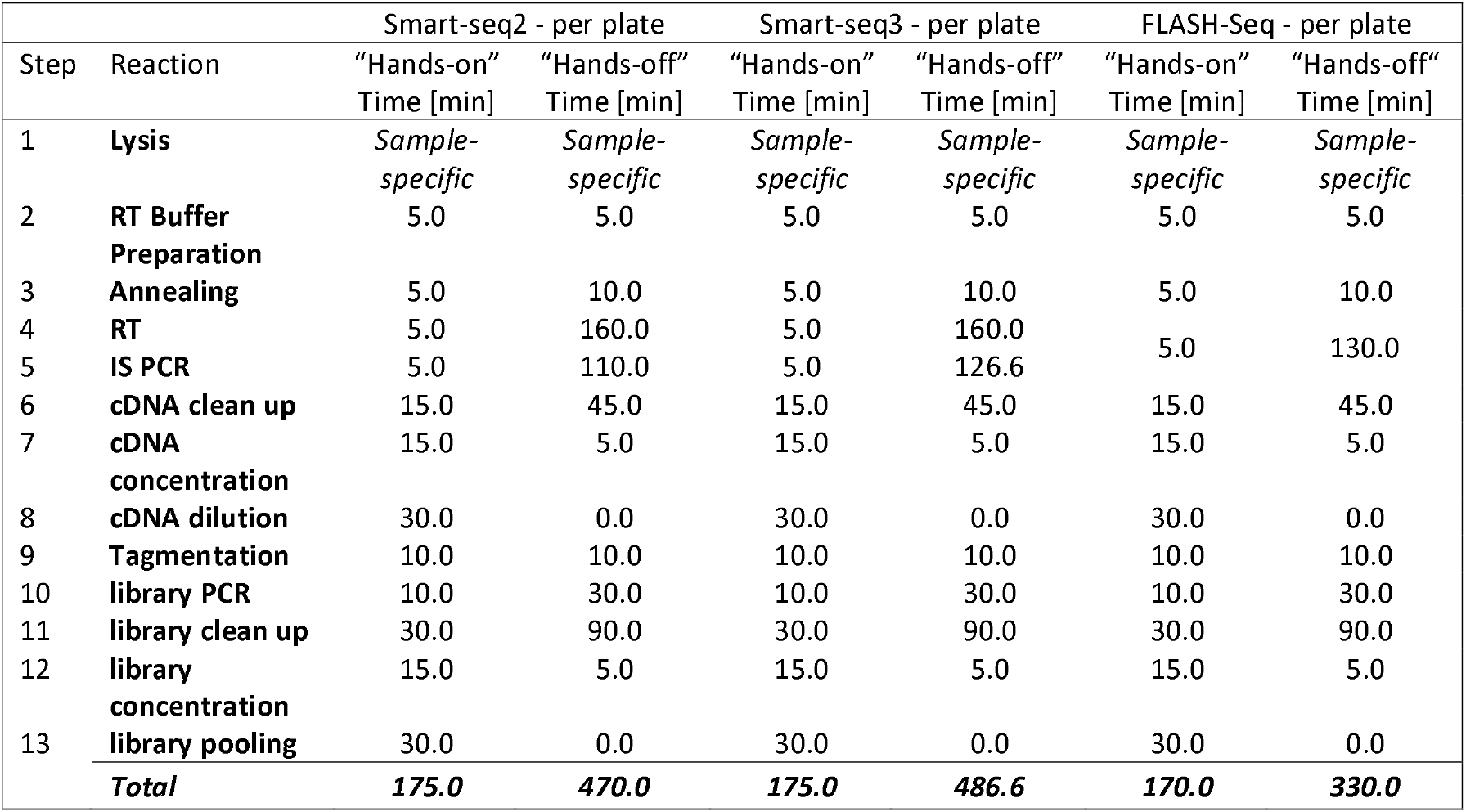
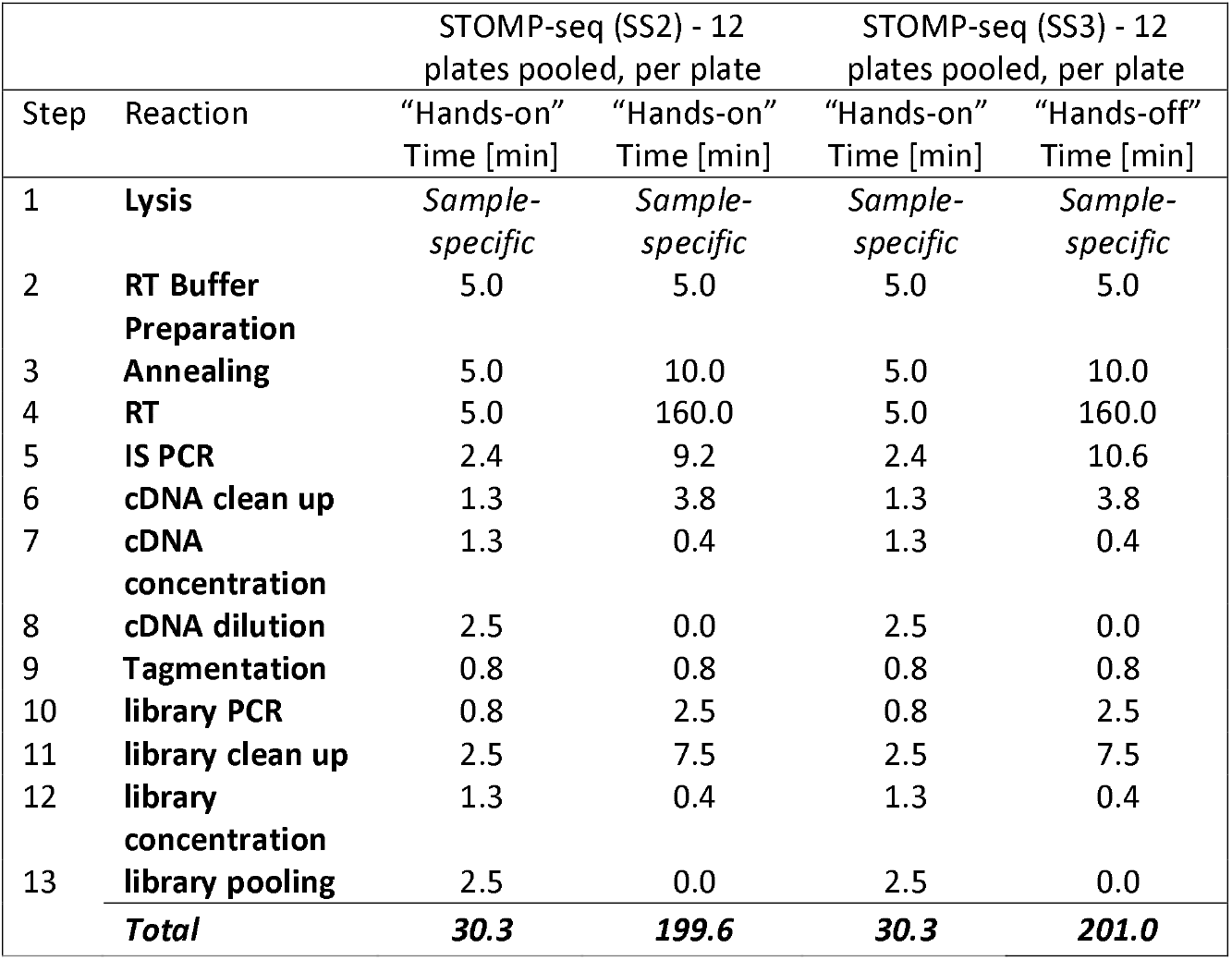
STOMP-seq protocol execution time compared to other SMART RNA-seq protocols, including reverse transcription (RT) cDNA amplification (IS) PCR, and sequencing library preparation (library) steps. FLASH-Seq execution times are estimated, performing equal library preparation steps from cDNA on as for Smart-seq2, Smart-seq3 and STOMP-seq.

**Supplementary Table 2.**
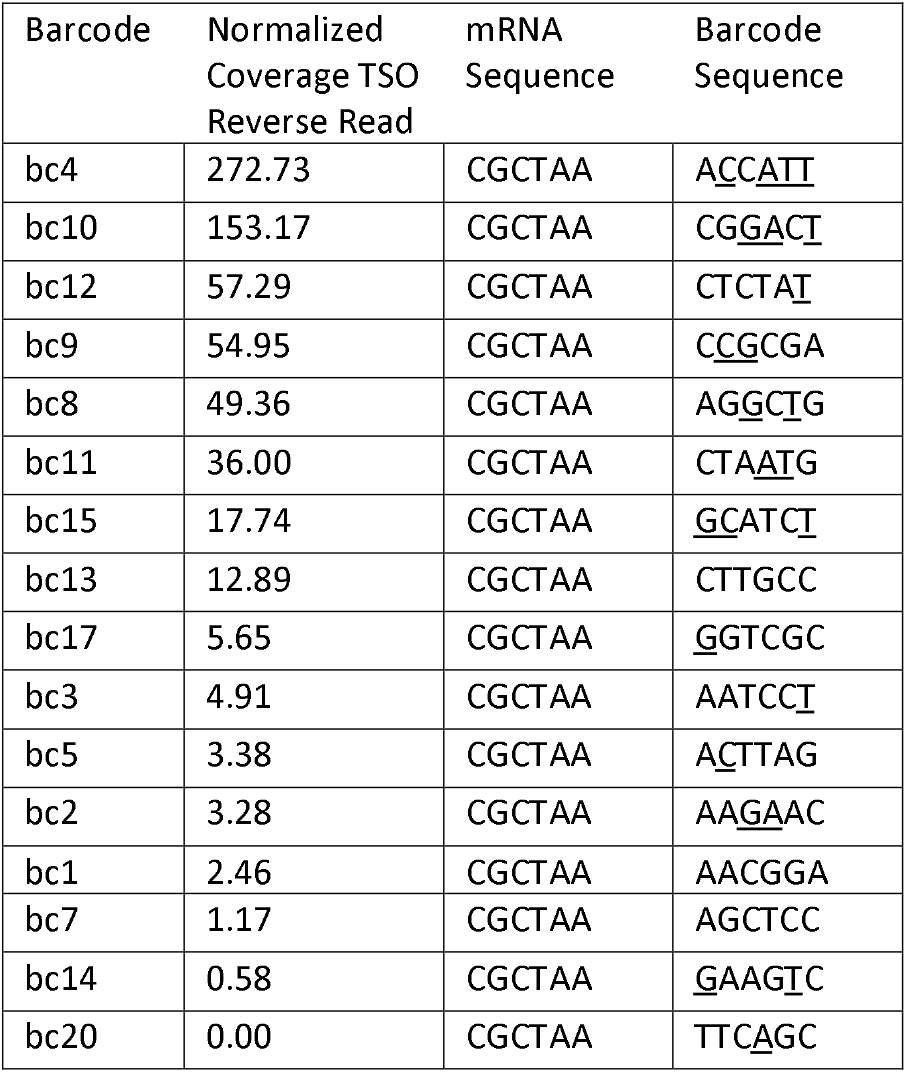
The influence of TSO barcode sequence on TSO mispriming. The number of reads derived from TSOs acting as PCR primers mapping to position 3020 of *vit-6* (Fig. S2e) for each barcode. Base complementarity between mRNA and barcode sequences is underlined.

**Supplementary Table 3.** STOMP-seq TSO barcode sequences evaluated in this study. (The 12 barcodes best-performing in Smart-seq2-based STOMP-seq are highlighted with asterisks.)

| Barcode name | Barcode sequence |
| --- | --- |
| bc1 | AACGGA |
| bc2* | AAGAAC |
| bc3 | AATCCT |
| bc4* | ACCATT |
| bc5* | ACTTAG |
| bc6 | AGAGAT |
| bc7* | AGCTCC |
| bc8 | AGGCTG |
| bc9* | CCGCGA |
| bc10* | CGGACT |
| bc11 | CTAATG |
| bc12* | CTCTAT |
| bc13 | CTTGCC |
| bc14* | GAAGTC |
| bc15* | GCATCT |
| bc16* | GGCAAG |
| bc17 | GGTCGC |
| bc18* | TCGTTC |
| bc19 | TCTGGT |
| bc20* | TTCAGC |
| bc21 | TTGGAG |
| bcSS2 | GTACAT |

**Supplementary Table 4.**
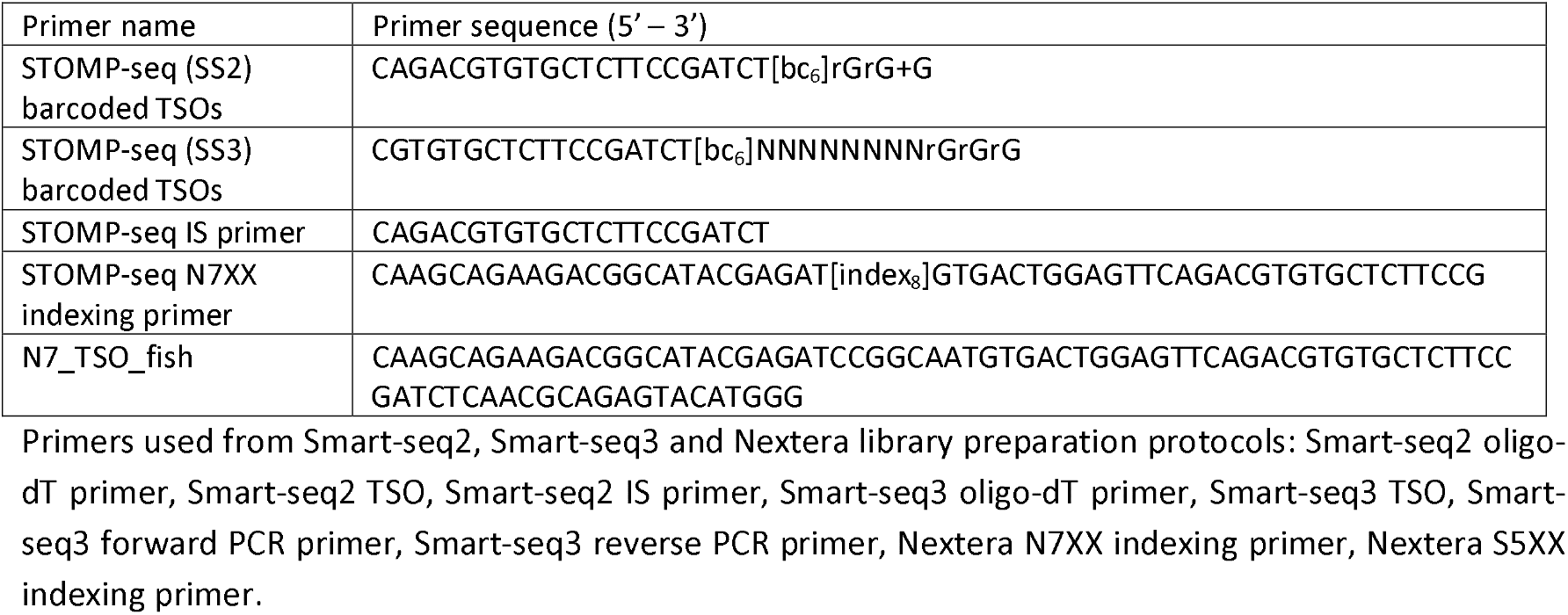
Primers and their sequences introduced in this study.

### Supplementary Figures

**Supplementary Figure 1.**
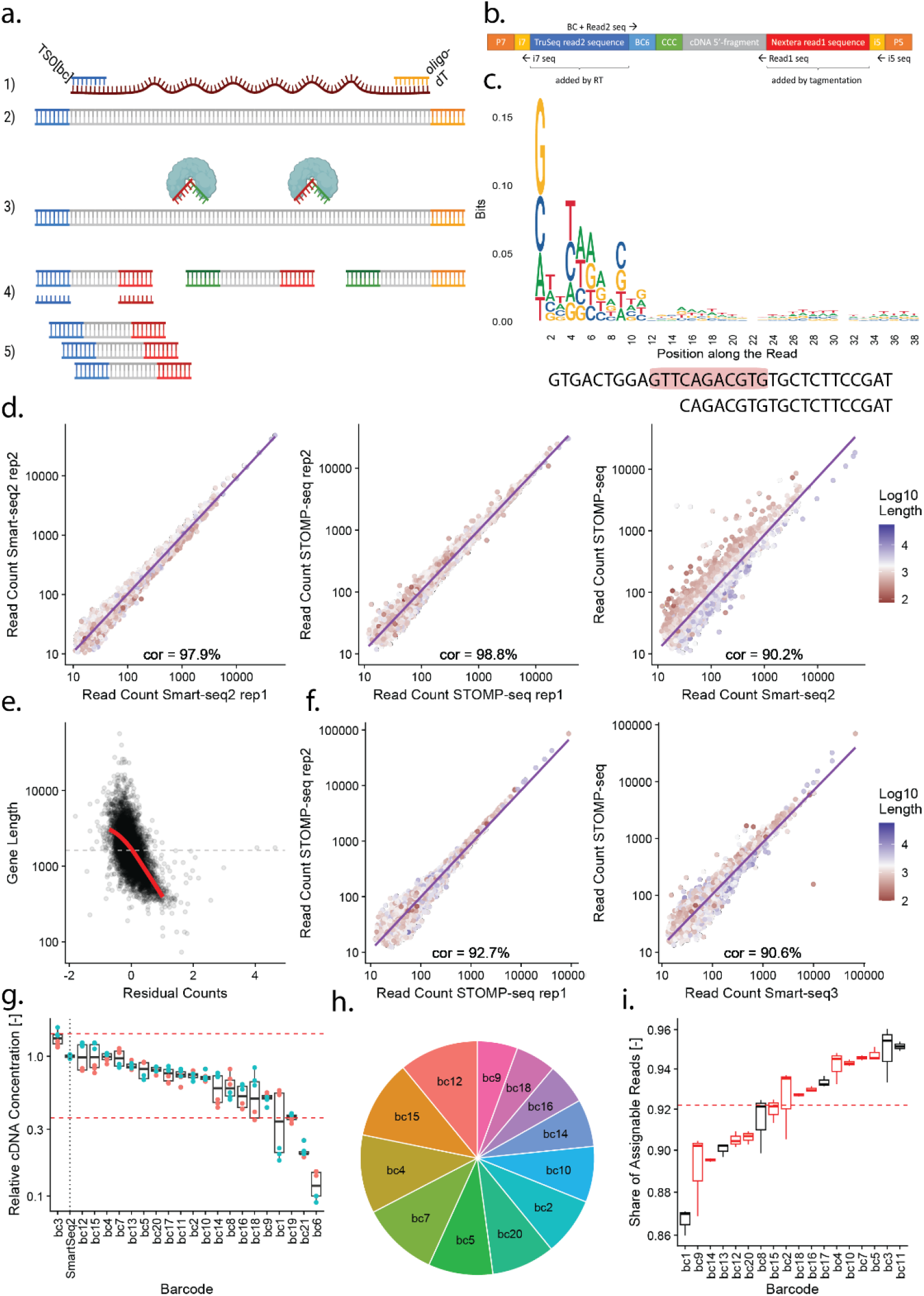
The STOMP-seq protocol enables a fast early-multiplexing strategy. **a**. Overview of the STOMP-seq protocol (*top*) with 1) cDNA generation and 2) amplification from poly-adenylated mRNA, 3) tagmentation with Tn5, loaded with Nextera primer sequences, and 4) cDNA 5’-fragment fishing and 5) amplification using the TruSeq sequence added through the TSO. **b**. Detailed composition of the resulting STOMP-seq sequencing fragment with sequences of interest highlighted. **c**. Deviation from randomness at the tagmentation site of tn5 (*top*). Full TruSeq read2 sequence, fragment with similarity to the Tn5 bias highlighted in red (*middle*). Truncated TruSeq read2 sequence with suppressed Tn5 bias used in STOMP-seq (*bottom*). **d**. The abundance of genes is consistent across two replicates of Smart-seq2 (*left*) and STOMP-seq (*middle*), with linear fit (*purple*). The average abundance of genes across 48 replicates processed with either Smart-seq2 or STOMP-seq (*right*). Each gene is colored by its coding sequence length, highlighting any bias; with linear fit (*purple*). **e**. Gene-length plotted against the residual difference in gene-expression between the two methods. **f**. The abundance of genes is consistent across two replicates of Smart-seq3-based STOMP-seq (*left*), with linear fit (*purple*). The average abundance of genes across 3 replicates processed with either Smart-seq3 or STOMP-seq (*right*). Each gene is colored by its coding sequence length, highlighting any bias; with linear fit (*purple*). **g**. cDNA yield of assessed barcoded TSOs compared to the cDNA yield achieved with Smart-seq2. Normalized technical triplicates of two biological replicates (*red, blue*). Black pointed line highlights the Smart-seq2 control. Red dotted lines mark the +-0.5x interval of the barcoded samples’ mean. **h**. Abundance of barcoded samples in a 12 fold multiplexed STOMP-seq cDNA library. **i**. The effect of STOMP-seq TSO barcodes on the fraction of reads containing an identifiable barcode, assessed in triplicate. Red dotted line indicates average across all barcodes, with the final twelve STOMP-seq barcodes chosen highlighted in red.

**Supplementary Figure 2.**
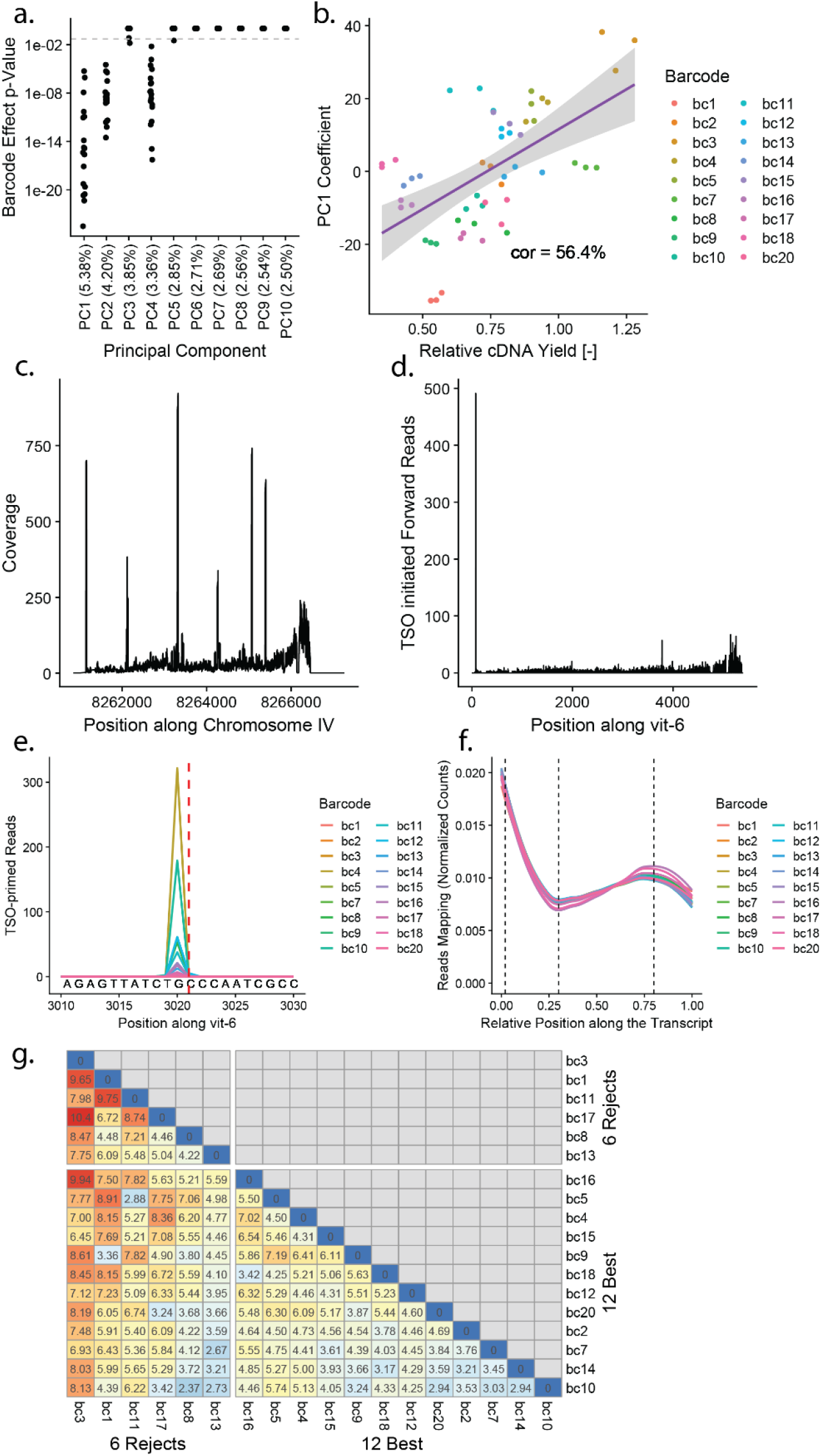
The effects of TSO barcode sequence on STOMP-seq cDNA libraries. **a**.The adjusted p-value of the barcode effect on the PC coefficient plotted for the first 10 PCs from PCA. **b**.Sample PC1 coefficients plotted against the respective obtained cDNA yield of barcoded samples, with linear model fit (*purple*). **c**. The read coverage of cDNA sequences adjacent to the Smart-seq2 TSO, mapping to the highly-expressed gene *vit-6*. **d**. The coverage of reads from cDNA sequences adjacent to the Smart-seq2 TSO, resulting from template switching at the 5’-end of *vit-6*. **e**. Visualization of the extend of TSO priming at position 3020 of *vit-6*, following three cytosines on the mRNA (*red dotted line*), colored by barcode. **f**. The relative read coverage averaged along the 1000 most highly-expressed genes, in libraries generated using different TSO barcode sequences (*colors*). Dotted lines indicate 5’, middle, and 3’ positions used to calculate coverage ratios. **g**. Heatmap of false positive differentially expressed genes inflation over false positive rates expected for p<.01, identified using triplicates of identical samples amplified using one barcode each. Comparisons between the final 12 chosen STOMP-seq barcode sequences are grouped together on the bottom right.

**Supplementary Figure 3.**
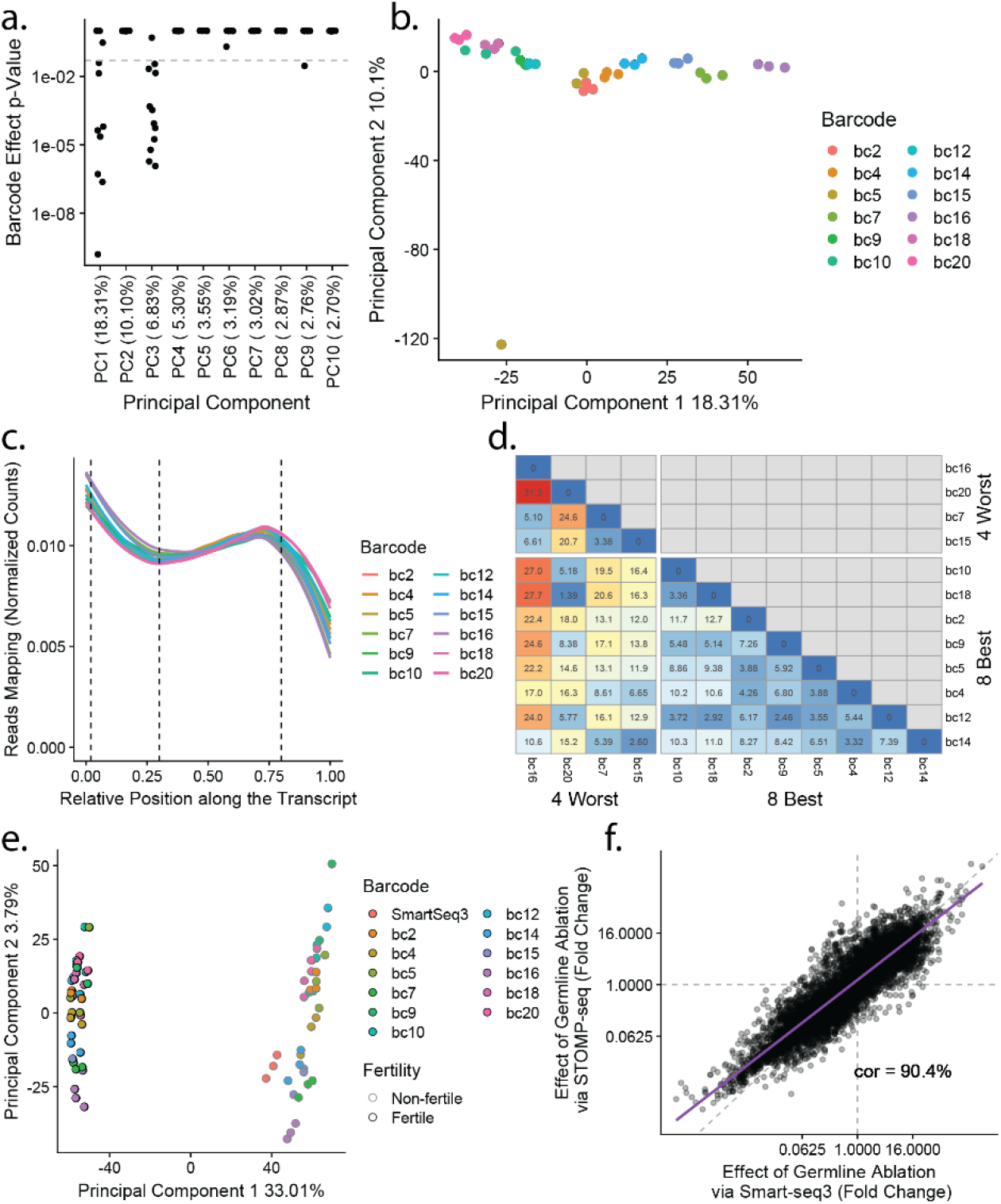
Evaluation of cDNA libraries generated with Smart-seq3-based STOMP-seq. **a**. The adjusted p-value of the barcode effect on the PC coefficient plotted for the first 10 PCs from PCA. **b**. Three replicates of Smart-seq3-based STOMP-seq were performed using each of 12 different TSO barcode sequences on identical aliquots taken from the same biological sample. Principal component analysis identifies two main axes of technical noise contributing 18.3% and 10.1% of all inter-sample variation, respectively. **c**. The relative read coverage averaged along the 1000 most highly-expressed genes, in libraries generated with Smart-seq3-based STOMP-seq using different TSO barcode sequences (*colors*). Dotted lines indicate 5’, middle, and 3’ positions used to calculate coverage ratios. **d**. Heatmap of false positive differentially expressed genes inflation over false positive rates expected for p<.01, identified using triplicates of identical samples amplified with Smart-seq3-based STOMP-seq using one barcode each. Comparisons between the 8 best STOMP-seq barcode sequences are grouped together on the bottom right. **e**. PCA of the transcriptomes of 3 intact (*black outline*) and 3 germline-ablated *glp-1(e2141*) (no outline) pools, processed either in STOMP-seq multiplexed pools (*colors*) or using Smart-seq3 (*red*). **f**. The fold-change effect of germline ablation on each gene’s expression, measured by Smart-seq3 (x-axis) and STOMP-seq (y axis), no significant differences were found between protocols at p<.01; with linear fit (*purple*).

**Supplementary Figure 4.**
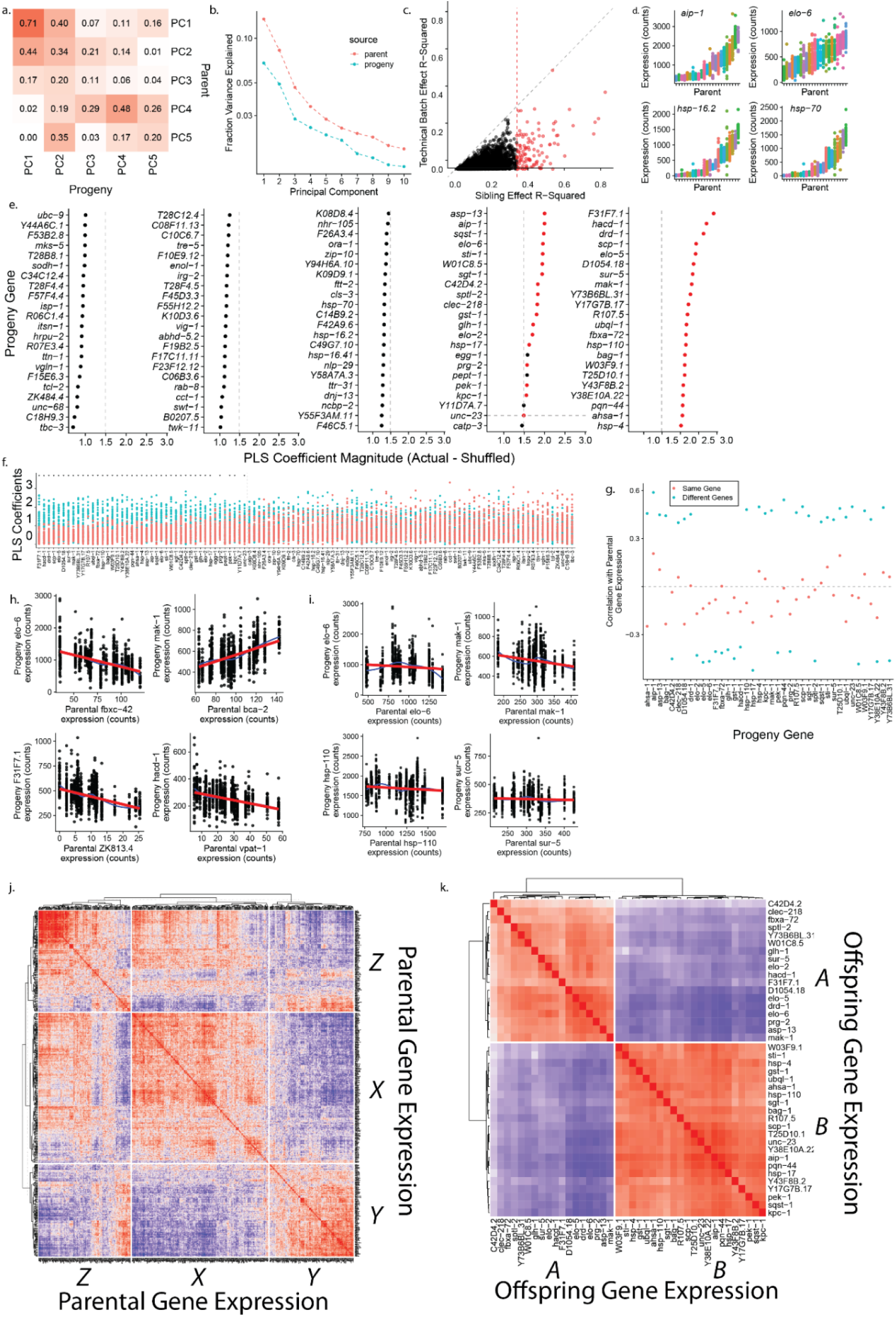
Using population-scale STOMP-seq to identify heritable gene-expression states. **a**. The correlation between the weights (loadings) of the first five principal components (PCs), calculated for all progeny (*x-axis*) vs all parents (*y-axis*). **b**. The fraction of variance explained by each PC in the respective populations. **c**. The fraction of gene-expression variance explained by sibling groups for each gene, compared to the fraction explained by shared technical groups (worm handling batch + sequencing batch). The cutoff of sibling effects at R^2^ > 0.4 is highlighted in red. **d**. Four example genes identified as showing greater similarity in expression between siblings (*shared colors*) than between cousins (*varied colors*). Each point represents the expression in a single individual. The same as in panel 2d, but here plotted using raw counts. **e**. The remaining PLS coefficients computed from real vs shuffled data. **f**. The PLS coefficients estimated using real parental data (*blue*) compared to shuffled parental data (*red*), i.e. the underlying data used to calculate panel e. Genes with statistically significantly higher coefficients in real data are decorated with an asterisk. **g**. For the set of 40 genes identified as having heritable expression levels, the Pearson correlation in expression was calculated between each gene (*x-axis*) and the most predictive gene in the parental transcriptome (*blue*). Separately, the correlation between the same gene (*x-axis*) in both progeny and parent (*red*) was calculated. **h**. Four example genes whose expression in progeny is well predicted by a single gene in the parents. **i**. Four example genes whose expression in progeny is well predicted by a single gene in the parents, but whose expression in progeny is not well predicted by the same gene’s expression in the parent. **j**. Within the parental population, hierarchical clustering identifies a more complex correlation pattern between genes in groups X, Y, and Z. k. Within the progeny population, hierarchical clustering reveals strong correlations between the genes in groups A and B.

